# Quantitative Proteomics Identifies Conserved Proteins and Altered Regulation of Mucin-16 in Low Grade Serous Ovarian Cancers

**DOI:** 10.1101/2024.04.29.591278

**Authors:** Christopher M. Tarney, Paulette Mhawech-Fauceglia, Jonathan D. Ogata, Julie Oliver, Tamara Abulez, Philip A. Branton, Saeid Movahedi-Lankarani, Brian L. Hood, Kelly A. Conrads, Kendal Rosalik, Kwong-Kwok Wong, David M. Gershenson, Sanghoon Lee, Anil K. Sood, Robert C. Bast, Kathleen M. Darcy, Neil T. Phippen, G. Larry Maxwell, Thomas P. Conrads, Nicholas W. Bateman

## Abstract

**Background:** Low-grade serous ovarian carcinoma (LGSOC) is a rare and largely chemoresistant subtype of epithelial ovarian cancer. Unlike treatment for high-grade serous ovarian cancer (HGSOC), management options for LGSOC patients are limited, in part, due to a lack of deep molecular characterization of this disease. To address this limitation, we aimed to define highly conserved proteome alterations in LGSOC by performing deep quantitative proteomic analysis of tumors collected from LGSOC (n=12) and HGSOC (n=24) patients or normal fallopian tube tissues (n=12) and validating proteins within two independent proteomic datasets of LGSOC (n=25) and HGSOC (n=49) tumors.

**Results:** Our efforts identified 275 protein alterations conserved between LGSOC and HGSOC tumors that exhibit high quantitative correlation between discovery and validation cohorts (Spearman Rho ≥ 0.82, P < 1E-4). Conserved proteins elevated in LGSOC tumors were enriched for pathways regulating cell adhesion and defective cellular apoptosis signaling and candidates mapping as putative drug targets included 5’-nucleotidase/ cluster of differentiation 73 (NT5E/CD73). We also identified MUC16 (CA125) as significantly elevated in LGSOC versus HGSOC tumors and confirmed this by immunohistochemistry analysis between three independent reviewers. We also find that MUC16 exhibits a more apical versus membrane-staining pattern in LGSOC tumors, suggesting unique regulation of MUC16 in this disease subtype.

**Conclusion:** Our efforts define highly conserved protein alterations distinguishing LGSOC from HGSOC tumors, including CD73, as well as the novel identification that MUC16 is elevated and exhibits more apical staining pattern in LGSOC tumor tissues. These findings deepen our molecular understanding of LGSOC and provide unique insights into highly conserved proteome alterations in LGSOC tumors.

## Introduction

Despite overall improvements in mortality, ovarian cancer is one of the most lethal gynecologic malignancies with fewer than half of women surviving five years beyond initial diagnosis^1^. In 2024, the American Cancer Society estimates that 19,680 women will be diagnosed with ovarian cancer and 12,740 (64%) will die from their disease^2^. Epithelial ovarian cancer is the most common histologic subtype of ovarian cancer and includes high-grade serous, low-grade serous, mucinous, clear cell, endometrioid, and carcinosarcoma. High-grade serous ovarian cancer (HGSOC) is the most common histologic subtype and is treated with a combination of cytoreductive surgery and platinum-taxane chemotherapy. Low-grade serous ovarian cancer (LGSOC) is responsible for 2-5% of ovarian cancer and is an overall challenging disease to treat given the high rate of chemoresistance of this disease compared to HGSOC^3^.

Historically, the hypothesis was that LGSOC ultimately progressed to HGSOC, but current opinion suggests these two disease processes areindependent^4^. From a molecular standpoint, both HGSOC and LGSOC vary by morphology and somatic mutations. HGSOC is associated with TP53 mutations while LGSOC often harbors mutations in Kirsten rat sarcoma virus (KRAS), the proto-oncogene B-Raf (BRAF), receptor tyrosine-protein kinase erbB-2 (ERBB2), beta-catenin (CTNNB1), and phosphatidylinositol 4,5-bisphosphate 3-kinase catalytic subunit alpha isoform (PIK3CA)^4^. However, co-existing high-grade and low-grade tumors often exhibit the same mutation and lack a TP53 mutation, suggesting there may be a stepwise, non-TP53 related pathogenic process^4^. More importantly, an understanding of genomic differences between these disease subtypes has also introduced new treatments for LGSOC patients ^5^.

Although the proteomic characterization of HGSOC is more widely described, there is currently limited proteomic data characterizing LGSOC tumors^6,7^. Some hypotheses suggest that LGSOC and HGSOC may originate from fallopian tube epithelium^3,8^, which provides a rationale for its incorporation in the proteomic evaluation of these tumors. As a result, we sought to perform a deep proteomic investigation evaluating tumors from patients diagnosed with LGSOC and HGSOC as well as normal fallopian tube tissues from women with benign disease to better understand the differences in molecular alterations between these tumors. We further aimed to define highly conserved proteome alterations between LGSOC and HGSOC tumors by integrating our discovery findings with previously described proteomic analyses of these tumor types^6,7,9^.

## Methods

### Patient cohort and tissue collections

Formalin-fixed, paraffin-embedded (FFPE) LGSOC and HGSOC tumor and normal fallopian tube tissues were obtained from Inova Fairfax Hospital (Falls Church, VA) under a less than minimal-risk WCG IRB-approved protocol #20122048 with waivers for consent and HIPAA authorization and evaluated under protocol #14-1679 with an exempt determination by the WCG IRB in accordance with the use of de-identified data under US Federal regulation 45 CFR 46.102(f). Benign fallopian tube tissues (FT) (n=12), as well as tumor tissues from LGSOC (n=12) and HGSOC (n=24) patients, were sectioned onto 8 µm polyethylene naphthalate membrane slides and hematoxylin-eosin stained. Equivalent areas of whole tissue from FT and tumor epithelium were collected using laser microdissection, avoiding areas of necrosis and hemorrhage, directly into 50 µL of Optima^TM^ LC/MS Grade water (Fisher Chemical) in microcentrifuge tubes and stored at - 80°C until sample preparation.

### Sample preparation for quantitative proteomic analysis

Protein from FFPE tissues was prepared for quantitative mass spectrometry (MS)-based proteomics as previously described^10^. Briefly, tissues were transferred to MicroTubes (Pressure BioSciences, Inc., Easton, MA, USA) including 20 µL of 100 mM TEAB (pH 8.0)/10% acetonitrile and underwent pressure-assisted trypsin digestion employing a barocycler (2320EXT, Pressure BioSciences, Inc.) and a heat-stable form of trypsin (SMART Trypsin, Thermo Fisher Scientific, Inc., Waltham, MA, USA). Peptide digest concentrations were determined using the bicinchoninic acid assay (BCA; Thermo Fisher Scientific, Inc.). Peptides were labeled with isobaric tandem mass tag (TMT) reagents according to the manufacturer’s instructions (TMT11^TM^ Isobaric Label Reagent Set, Thermo Fisher Scientific). Sample multiplexes were reversed-phase fractionated (basic pH) on a 1260 Infinity II offline liquid chromatography system (Agilent Technologies, Inc., Santa Clara, CA, USA) into 96 fractions using a linear gradient of acetonitrile (0.69% min^-1^) followed by concatenation into 24 pooled fractions. Pooled fractions were lyophilized in a Speed-Vac and stored at -20 °C until resuspended in 25 mM NH_4_HCO_3_ and analyzed by liquid chromatography-tandem mass spectrometry (LC-MS/MS).

### LC-MS/MS Analysis

Peptide identifications and quantitation for TMT multiplexes were performed as recently described^9^. Liquid chromatography-tandem mass spectrometry (LC-MS/MS) analyses of TMT11 multiplexes were performed on a nanoflow high-performance LC system (EASY-nLC 1200, Thermo Fisher Scientific, Inc.) coupled online with an Orbitrap mass spectrometer (Q Exactive HF-X, Thermo Fisher Scientific, Inc.). Samples were loaded on a reversed-phase trap column (Acclaim^TM^ PepMap^TM^ 100 Å, C-18, 20 mm length, nanoViper Trap column, Thermo Fisher Scientific, Inc.) and eluted on a heated (50 °C) reversed-phase analytical column (Acclaim^TM^ PepMap^TM^ RSLC C-18, 2 μm, 100 Å, 75 μm × 500 mm, nanoViper, Thermo Fisher Scientific, Inc.) by developing a linear gradient from 2% mobile phase A (2% acetonitrile, 0.1% formic acid) to 32% mobile phase B (95% acetonitrile, 0.1% formic acid) over 120 min at a constant flow rate of 250 nL/min. Full scan mass spectra (MS) were acquired using a mass range of *m/z* 400-1600, followed by selection of the top 12 most intense molecular ions in each MS scan for high-energy collisional dissociation (HCD). Instrument parameters were as follows: Full MS: AGC, 3 × 10e6; resolution, 60 k; S-Lens RF, 40%; max IT, 45 ms; MS2: AGC, 10e5; resolution, 45 k; max IT, 95 ms; quadrupole isolation, 1.0 *m/z*; isolation offset, 0.2 *m/z*; NCE, 34; fixed first mass, 100 *m/z*; intensity threshold, 2 × 10e5; charge state, 2–4; dynamic exclusion, 20 s with 10 ppm tolerance, TMT optimization. Global protein-level abundances were generated from peptide spectral matches identified by searching .raw data files against a publicly available, non-redundant human proteome database (http://www.uniprot.org/, SwissProt, Homo sapiens, downloaded 12-01-2017 using Mascot (Matrix Science, v2.6.0), Proteome Discoverer (v2.2.0.388, Thermo Fisher Scientific, Inc.), and in-house tools using identical parameters as previously described^9^.

### Bioinformatic Analysis

Differential analyses of TMT-11 data matrixes were performed using the LIMMA package (version 3.8)^11^ in R (version 3.6) and Mann-Whitney U rank sum testing in MedCalc (version 19.0.3). Pathway analysis was performed using MSigDB and GSEA^12^ considering hallmark and canonical pathways using default enrichment settings, i.e. significantly enriched pathways satisfied a false discovery rate < 0.05 or STRING (https://string-db.org/) using default parameters for Homo Sapiens. Visualizations were generated using SRplot^13^, GraphPad Prism (v.10.0.3), or MedCalc. Global proteome data for LGSOC and HGSOC validation cohorts were previously reported^6,7,9^ and HGSOC data^9^ prioritized for this analysis was focused on tumors where tissue source site was designated as primary.

### Immunohistochemical analysis of mucin-16 (MUC16)/ cancer antigen 125 (CA125)

LGSOC and HGSOC tissues were sectioned onto charged glass slides, fixed with 100% methanol, incubated at 5 min at ambient temperature then rinsed with phosphate-buffered saline (PBS). Tissue was permeabilized by incubating with 0.05% triton-PBS for 15 min and 2.5% normal goat serum. The slides were incubated overnight at 4 °C with anti-CA125 mouse monoclonal antibody (Clone M11, Agilent Dako, Santa Clara, CA, USA 1:1000). Dako’s Envision diaminobenzidine (DAB) detection system was used to label bound antibody complexes. Normal lung tissue was used as the positive tissue control. Following detection, slides were counterstained with hematoxylin then dehydrated and cover slipped. Stained tissue sections were scanned on an Aperio ScanScope XT slide scanner (Leica Microsystems, Wetzlar, Germany). Digital images underwent expert pathology review (P.M.F., P.A.B., S.M.L.) and a Cohen’s kappa statistic was calculated to assess inter-rater variability of MUC16-positive tumor cell reviews categorized as quintiles using the irr package (version 0.84.1). Odds ratio analysis was performed by comparing tumors classified as exhibiting a “Apical” or “Apical/ Membranous” staining pattern designations by expert pathology review using MedCalc (version 20.109).

## Results

### Quantitative proteomic analyses of whole tissue collections from normal fallopian tube as well as low and high-grade serous ovarian cancers

We performed multiplexed, quantitative proteomic analyses of whole tissue equivalents collected from low-grade (LGSOC, n=12) or high-grade serous ovarian cancer (HGSOC, n=24) tumors and fallopian tube tissues (FT, n=12) from women with benign disease (Figure 1, Table 1, Supplementary Table 1). Global proteomic analyses resulted in the quantitation of 7,043 proteins across all patient samples (Supplementary Table 2). Principal component analysis of top variably abundant proteins quantified across all patient samples explained 37.1% (PC1) and 8.9% (PC2) of the variance between FT, LGSOC, and HGSOC tissue collections (Figure 2A). Differential analysis of proteins quantified between HGSOC or LGSOC and FT collections identified both co-altered as well as unique protein alterations between these comparisons (Figure 2B, Supplementary Table 3 and 4).

**Figure 1:**
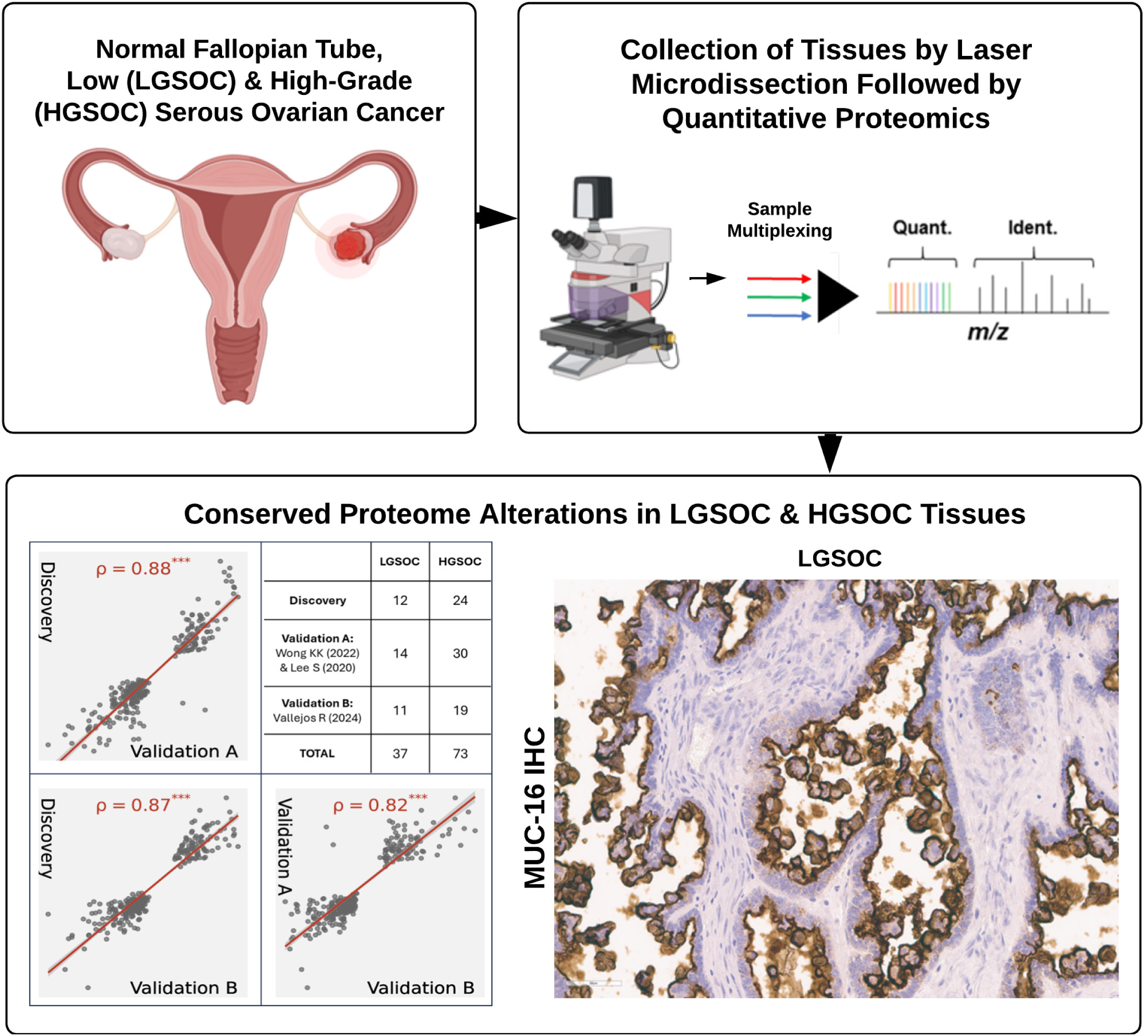
Workflow supporting quantitative proteomic analyses and validation of proteome alterations in low (LGSOC) and high-grade (HGSOC) serous ovarian cancers. Whole tissue collections from benign fallopian tube, LGSOC, and HGSOC tissues were performed using laser microdissection. Tissue collections underwent multiplexed, quantitative proteomic analysis using isobaric tagging, i.e. tandem mass tags (TMT-11), followed by liquid chromatography, tandem mass spectrometry (LC-MS/MS) analysis on an Orbitrap mass spectrometer. Proteome alterations identified in discovery cohorts of LGSOC and HGSOC tissues were validated using global proteome data for independent cohorts of LGSOC (n=25) and HGSOC (n=49) tumors^6,7,9^ and included the finding that mucin-16 (MUC16) is elevated in LGSOC tumors which was further verified by immunohistochemistry analysis of LGSOC and HGSOC tissues by three independent reviewers.

**Figure 2:**
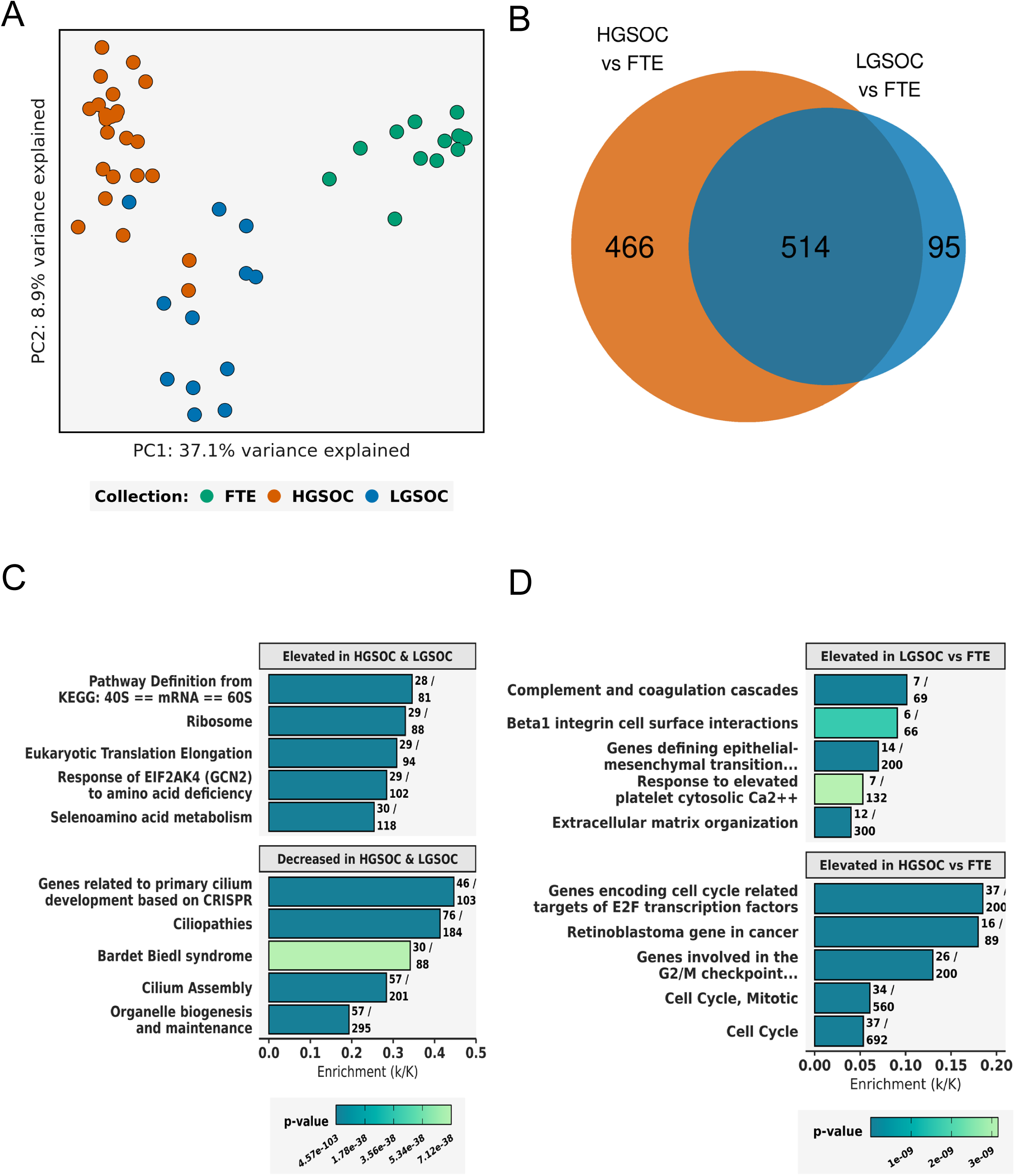
Differential proteomic analysis of whole tissue collections from normal fallopian tube with low (LGSOC) and high-grade (HGSOC) serous ovarian cancers. A: Principal component analysis (PCA) of the top 500 most variably abundant proteins quantified across samples; proteins prioritized based on median absolute deviation. B: Comparison of proteins significantly altered between HGSOC or LGSOC and benign fallopian tube collections; proteins reflect LIMMA adjusted p-value < 0.05, fold-change ± 2.0. C: Pathway analysis of proteins co-altered between HGSOC and LGSOC vs benign fallopian tube collections; top hallmark and canonical pathways (gsea-msigdb.org) enriched among proteins elevated (84) or decreased (457) in HGSOC and LGSOC versus benign fallopian tube collections. D: Pathway analysis of proteins uniquely elevated in HGSOC or LGSOC vs normal fallopian tube collections; top hallmark and canonical pathways (gsea-msigdb.org) enriched among proteins elevated in HGSOC (300) or LGSOC (158) tumors versus benign fallopian tube collections.

**Table 1:** Patient and tumor clinicopathological characteristics.

| Characteristic | High Grade Serous<br>Ovarian Cancer<br>(N = 24) | Low Grade Serous<br>Ovarian Cancer<br>(N = 12) | Benign<br>Fallopian Tube<br>(N = 12) |
| --- | --- | --- | --- |
| Mean age (range) –yr | 58.1 (30-90) | 52.0 (30-73) | 45.2 (41-49) |
| Menopausal status – no. (%) |  |  |  |
| Premenopausal | 5 (20.8) | 6 (50.0) | 12 (100.0) |
| Postmenopausal | 19 (79.2) | 6 (50.0) | 0 (0.0) |
| Mean BMI (range) | 25.6 (22.5 – 30) | 25.4 (20.7 – 30.8) | 30.1 (19 – 44.1) |
| Race – no. (%) |  |  |  |
| White | 13 (54.2) | 11 (91.7) | 4 (33.3) |
| Black | 7 (29.2) | 0 (0.0) | 4 (33.3) |
| Asian | 4 (16.7) | 1 (8.3) | 2 (16.7) |
| Hispanic | 0 (0.0) | 0 (0.0) | 2 (16.7) |
| FIGO stage – no. (%) |  |  |  |
| I or II | 2 (8.3) | 3 (25.0) | - |
| IIIA | 3 (12.5) | 4 (33.3) | - |
| IIIB | 6 (25.0) | 1 (8.4) | - |
| IIIC | 7 (29.2) | 4 (33.3) | - |
| IV | 6 (25.0) | 0 (0.0) | - |
| Debulking |  |  |  |
| RD0 | 12 (50.0) | 7 (58.3) | - |
| RD1 | 9 (37.5) | 4 (33.3) | - |
| Suboptimal | 2 (8.3) | 1 (8.4) | - |
| Unknown | 1 (4.2) | 0 (0.0) | - |
| Platinum Sensitivity – no (%) |  |  |  |
| Sensitive | 14 (58.3) | 8 (66.6) | - |
| Resistant | 4 (16.7) | 1 (8.4) | - |
| Unknown | 6 (25.0) | 3 (25.0) | - |
| Mean CA125 (range) | 4917.1 (329 – 50,000) | 672.7 (70 – 1576) | - |

Pathway analysis of co-altered proteins elevated in HGSOC and LGSOC compared to FT identified enrichment of top pathways correlating with regulation of protein translation (i.e., 40S/ 60S ribosome and eukaryotic translation elongation) as well as protein metabolism (i.e., response of EIF2AK4 to amino acid deficiency and selenoamino acid metabolism) in tumor samples (Figure 2C, Supplementary Table 5). Pathways enriched among proteins elevated in FT tissues largely correlated with regulation of cilium development as well as organelle biogenesis Figure 2C, Supplementary Table 5). Proteins uniquely elevated in HGSOC vs FT identified pathways largely associated with cell cycle regulation while those enriched for proteins uniquely elevated in LGSOC vs FT correlated with pathways regulating epithelial to mesenchymal transition and hematopoietic cell-related signaling (Figure 2D, Supplementary Table 6).

### Characterization and validation of proteome alterations between low and high-grade serous ovarian cancers

Our discovery cohort of LGSOC and HGSOC tumors was compared with global proteome data of tumor tissues collected from two independent cohorts of women diagnosed with LGSOC^6,7^ (n=25) or HGSOC^7,9^ (n=49) (Figure 3A). Differential analysis of proteins quantified in LGSOC and HGSOC tumors identified 785 proteins as significantly altered between these tumor types (Figure 3B). Top protein alterations included proliferation marker protein Ki-67 (MKI67) elevated in HGSOC and multiple collagen isoforms in LGSOC tumors (Figure 3B, Supplementary Table 7). Comparison of these protein alterations across discovery and validation cohorts identified 275 significantly co-altered proteins between these disease subtypes that further exhibit high quantitative correlation (Figure 3C) with two proteins, dihydropyrimidine dehydrogenase [NADP(+)] (DPYD) and Fructose-1,6-bisphosphatase 1 (FBP1), exhibiting discordant abundance between cohorts (Supplementary Table 8). Pathway analysis of these features showed proteins elevated in LGSOC tumors were enriched for pathways regulating intercellular adhesion and defective cellular apoptosis signaling while proteins elevated in HGSOC tumors showed enrichment of pathways largely regulating DNA replication and cell cycle (Figure 3D3D, Supplementary Table 9). We further investigated proteins that are putative drug targets in LGSOC and HGSOC tumors and identified 27 features (Supplementary Table 8). Putative drug targets exhibiting the largest abundance differences between these tumor types included Annexin A1 (ANXA1) and 5’-nucleotidase/ cluster of differentiation 73 (NT5E/CD73) as elevated in LGSOC tumors and DNA topoisomerase 2-alpha (TOP2A) as well as cyclin-dependent kinase 1 (CDK1) as elevated in HGSOC tumors.

**Figure 3:**
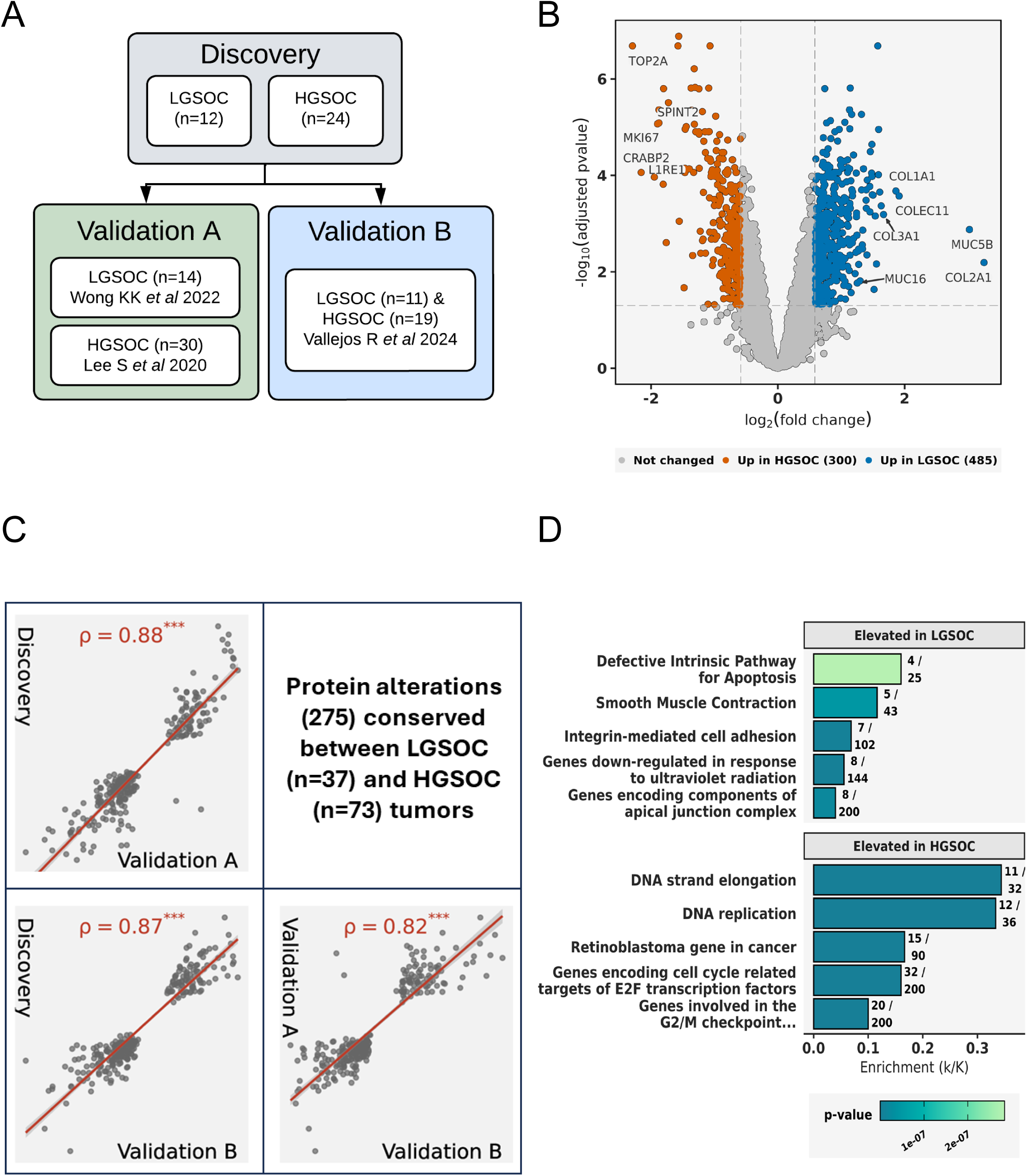
Differential proteomic analysis of low (LGSOC) and high-grade (HGSOC) serous ovarian cancers and validation of proteome alterations in independent LGSOC and HGSOC cohorts. A: Workflow for the prioritization of conserved protein alterations between LGSOC and HGSOC tumors in discovery and validation cohorts^6,7,9^. B: Volcano plot reflects proteins significantly altered between LGSOC and HGSOC tumors in discovery cohort; proteins reflect LIMMA adjusted p-value < 0.05, fold-change ± 1.5. C: Proteins co-altered between LGSOC and HGSOC tumors in discovery and validation cohorts; 275 co-altered proteins passing an adjusted p-value < 0.05. D: Pathway analysis of proteins co-altered between LGSOC and HGSOC tumors in discovery and validation cohorts; top hallmark and canonical pathways (gsea-msigdb.org) enriched among proteins elevated (93) or decreased (180) between LGSOC and HGSOC tumors.

We also identified mucin-16 (MUC-16), also known as cancer antigen 125 (CA125), as significantly elevated in LGSOC versus HGSOC tumors passing false-discovery rate (FDR) < 0.05 expectations in discovery and validation A cohorts (Figure 4A) and exhibiting concordant abundance in validation B cohort as elevated in LGSOC versus HGSOC tumors with an FDR = 0.052. Circulating MUC16/CA125 levels are assessed as part of routine diagnosis and disease monitoring in the clinical management of ovarian cancer^3,8^. To uncover whether other putatively conserved protein candidates elevated in LGSOC tumors are predicted to interact with MUC16, we investigated highly conserved protein alterations using the STRING protein interaction network (string-db.org) and identified nesprin-1 (SYNE1) as significantly elevated in LGSOC tumors (Supplementary Table 8). Considering the importance of MUC16 in ovarian cancer disease biology, we were motivated to investigate this protein alteration in LGSOC and HGSOC tissues further.

**Figure 4:**
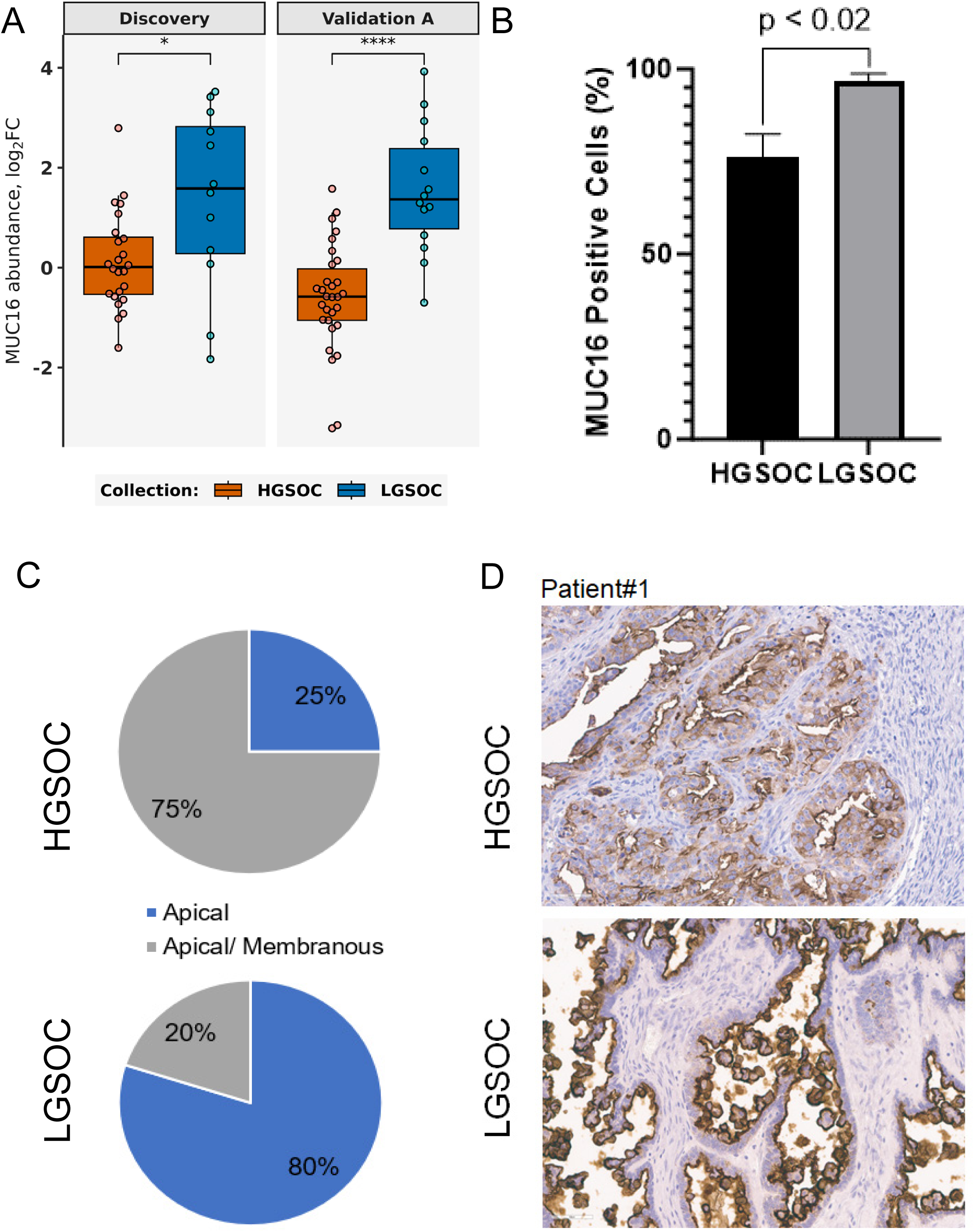
Immunohistochemical analysis of Mucin-16 (MUC16) in low (LGSOC) and high-grade (HGSOC) serous ovarian cancers. A: Mucin-16 (MUC16) protein abundance in LGSOC and HGSOC tumors from discovery or validation A cohorts; * = LIMMA adjusted p-value < 0.05, **** = LIMMA adjusted p-value < 1E-4. MUC16 was significantly elevated in LGSOC tumors in Validation B cohort, adjusted p-value= 0.052 (not shown). B: Comparison of cells stained positive for MUC16 following immunohistochemical analysis of LGSOC (n=10) and HGSOC (n=24) tissues; data and p-value reflects Mann Whitney Rank Sum testing comparing “PATH1” reviews (Supplementary Table 10). MUC16 confirmed elevated in LGSOC tumors by three independent reviewers (Cohen’s kappa = 0.462, P < 1E-4, Supplementary Table 10). C: Comparison of MUC16 staining patterns in LGSOC (n=10) and HGSOC (n=24) tissues identified by “PATH1”; tumor cell populations were identified to exhibit predominantly apical or apical-membranous MUC16 staining and predominantly apical staining in LGSOC tumors was confirmed by three independent reviewers (Fisher’s Exact P ≤ 0.034, Supplementary Table 10). D: MUC16 immunohistochemistry stains in tissue sections from one representative LGSOC and HGSOC patient tumor.

### Mucin-16 (MUC16)/ cancer antigen 125 (CA125) is elevated and exhibits a predominantly apical staining pattern in low versus high-grade serous ovarian cancers

To further investigate elevation of MUC16 protein as a putatively conserved protein alteration in LGSOC versus HGSOC tumors, we performed immunohistochemistry (IHC) analysis of tissues for a subset of LGSOC (n=11) and HGSOC (n=24) tumors characterized within our discovery cohort. A comparison of tumor cells stained positive for MUC16 between LGSOC and HGSOC tumors showed HGSOC tumors overall have fewer MUC16-positive cells than LGSOC cells (Figure 4B, Supplementary Table 10), a finding confirmed by three independent reviewers (Cohen’s kappa = 0.462, P < 1E-4, Supplementary Table 10). Further analysis of MUC16 staining identified discrete localization patterns correlating within disease subtypes, where HGSOCs exhibited a more membranous or combined apical and membranous staining pattern while LGSOCs exhibited a predominantly apical staining pattern for MUC16 (Figure 4C). The increased apical staining pattern in LGSOC tumors was also confirmed by three independent reviewers (Fisher’s Exact P ≤ 0.034, Supplementary Table 10). We further include MUC16 IHC images for a representative HGSOC case demonstrating apical and membranous staining, and an LGSOC case exhibiting a predominant and intense apical membrane staining pattern (Figure 4D).

## Discussion

Our deep proteomic analysis of benign fallopian tube tissues as well as low (LGSOC) and high (HGSOC) grade serous ovarian cancers shows these samples can be stratified by variably abundant protein alterations suggesting distinct proteome abundance profiles are characteristic of these tissue types. Differential analysis of tumor (LGSOC and HGSOC) and benign fallopian tube tissue identified that proteins elevated in fallopian tubes were largely enriched for pathways regulating cilium development. Recent evidence from single cell studies has identified several subpopulations of epithelial cell types exist within benign fallopian tube, including ciliated cell types^14,15^. This cell population is considered to reflect a more differentiated subpopulation of fallopian tube epithelial cells emerging from non-ciliated, secretory epithelial cell types^14,15^. The loss of proteins regulating cilium development that we observe in LGSOC and HGSOC tissues is consistent with evidence suggesting secretory epithelial cell types often exhibit greater neoplastic potential than other cell types in fallopian tube epithelium^14^. We also find that proteins elevated in both LGSOC and HGSOC tumors exhibit enrichment of pathways regulating protein translation, specifically ribosome and translational elongation, as well as amino acid metabolism in comparison with benign fallopian tube. Increased ribosome biogenesis and protein translation has been described as a characteristic of ovarian cancer cells and as a possible therapeutic vulnerability in both LGSOC and HGSOC^16,17^. Proteins uniquely elevated in HGSOC tumors compared to benign fallopian tube were largely enriched for pathways regulating cell cycle and mitosis, which is consistent with the high-grade phenotype of this disease^18^. Proteins uniquely elevated in LGSOC were enriched for pathways regulating epithelial to mesenchymal transition including extracellular matrix organization, beta1 integrin cell surface interactions, and hematopoietic cell-related signaling, i.e. complement and coagulation. Beta1 integrin signaling has been implicated in regulating migration of ovarian cancer cells in the context of interactions with extracellular matrix proteins^19^. Our comparison with fallopian tube tissues has identified proteome alterations largely consistent with the malignant phenotypes of LGSOC and HGSOC tumors in comparison to benign fallopian tube.

Comparison of LGSOC and HGSOC revealed marked protein alterations between these disease subtypes, including elevation of proteins correlating with proliferation and DNA replication in HGSOC. We also observed elevation of diverse extracellular matrix proteins in LGSOC tumors, including multiple collagen isoforms which is noteworthy considering the enrichment of pathways regulating intercellular cell adhesion^20^ we also observe in these tumors. Recent global proteome analysis of tumors collected from women diagnosed with LGSOC or HGSOC have been described by our team^6,9^ and others^7^, and we were motivated to compare observed proteome alterations between LGSOC and HGSOC tumors from this study with these independent cohorts to identify and validate proteome alterations that are highly conserved between these disease subtypes. This comparison resulted in the identification of highly conserved protein alterations exhibiting high quantitative correlation between LGSOC and HGSOC tumors (Spearman Rho ≥ 0.82). These features demonstrated that proteins elevated in HGSOC tumors correlate with enrichment of pathways regulating cell cycle and cell mitosis signaling, while those elevated in LGSOC tumors correlated with pathways regulating intercellular adhesion and defective cellular apoptosis. We also investigated putative drug targets exhibiting large differences in abundance between LGSOC and HGSOC tumors and, among top targets, we identify cluster of differentiation 73 (NT5E/CD73) as elevated in LGSOC versus HGSOC tumors. CD73 has been shown to participate in an immune-related checkpoint pathway that promotes conversion of exogenous adenosine triphosphate to adenosine by tumor cells which can suppress immune cell function ^21,22^. CD73 expression has been shown to correlate with poor disease outcome in HGSOC and to be markedly elevated in HGSOC tumors that are enriched for a stromal-expression signature also correlated with poor disease outcome^23^. Recent efforts investigating the feasibility to target CD73 with a monoclonal antibody, i.e. Oleclumab, and the immune checkpoint inhibitor, i.e. Durvalumab, in recurrent ovarian cancer have found this combination to be well tolerated^22^. LGSOC has been recently described as exhibiting an immunosuppressive tumor microenvironment, particularly in the absence of mutations in the proto-oncogene B-Raf (BRAF) or Kirsten rat sarcoma virus (KRAS)^24^. Our findings that CD73 is significantly elevated in LGSOC versus HGSOC tumors warrants additional investigation of the role of this protein in regulating the immune microenvironment of LGSOC tumors and will be the focus of future research.

Additional investigation of putatively conserved protein alterations between LGSOC and HGSOC tumors identified mucin-16 (MUC16)/ cancer antigen 125 (CA125) as significantly elevated in LGSOC versus HGSOC tumor tissues. MUC16 is a transmembrane glycoprotein that is highly abundant in the female reproductive tract and can be actively shed from ovarian cancer cells into blood plasma^8^. Circulating MUC16/ CA125 protein levels thus represents a measurable biomarker that can assess evidence of disease and is a facet of routine diagnosis and disease monitoring in the clinical management of ovarian cancer^3,8^. Comparison of the putative MUC16 interactome and highly conserved protein alterations observed between LGSOC and HGSOC tumors identified nesprin-1 (SYNE1) as elevated in LGSOC tumors. SYNE1 and MUC16 have been observed as commonly mutated gene targets in stomach cancer and tumors with mutations in these genes have been shown to exhibit decreased immune cells as well as to correlate with lower response to immunotherapy^25^. Shedding of MUC16 has been shown to occur as a product of proteolytic cleavage of the carboxy-terminal ectodomain in the acidifying Golgi/ post-Golgi^26^, possibly by membrane-type 1 matrix metalloproteinase^27^ followed by release of the N-terminal CA125 region^27^ into interstitial fluid and ultimately into circulating blood^8^. As we quantify membrane-type 1 matrix metalloproteinase (MT1-MMP/ MMP-14/ P50281) protein in our discovery cohort, we compared levels between LGSOC and HGSOC tumors, but we do not observe this protein as significantly altered between these disease subtypes (MWU p = 0.48). Recent evidence investigating mechanisms regulating cell surface retention of MUC16 have implicated N-glycosylation mediated by N-acetylglucosaminyltransferase I (MGAT1) which has been shown to promote increased retention of MUC16 at the cell surface^28^. This study further identified that N-glycosylation of MUC16 promotes interaction with the protein galectin-3 (LGALS3) at the cell surface^28^. Notably, we identify LGALS3 as significantly elevated in LGSOC versus HGSOC tumor tissues (p<0.05 across all cohorts assessed) and, although we quantify MGAT1, we see this protein is significantly elevated in HGSOC tumors in our discovery cohort (Supplementary Table 7), but we do not see this protein as significantly altered between LGSOC versus HGSOC tissues in validation cohorts. If we consider this evidence in the context of our findings, this suggests LGSOC tumor cells perhaps exhibit increased retention of MUC16 at the cell surface. Circulating CA125 levels have been measured in both LGSOC and HGSOC patients, although it has been noted that these levels are more commonly lower in LGSOC patients^3,29^. We also find that circulating CA125 levels were lower in our LGSOC versus HGSOC patients (Table 1), providing further evidence in support of this hypothesis. A possible mechanism underlying altered dynamics of MUC16 retention may be due to differential N-glycosylation of MUC16 in LGSOC tumor cells, possibly due to altered enzymatic activity of MGAT1. Alternatively, differences in MUC16 expression and shedding could relate to greater cleavage of the extracellular domain in HGSOC, accounting for the greater shedding of CA125 and decreased membrane expression. Future studies will investigate these hypotheses, including the comparison of cleaved versus non-cleaved MUC16 levels in LGSOC and HGSOC tumors.

We also observe that LGSOC tumor cells exhibit a predominantly apical staining pattern for MUC16 in comparison to HGSOC tumor cells. Similar findings were observed in murine ovarian carcinomas where more intense apical expression of MUC16 was observed in grade 1 tumors^30^. Recent evidence has shown that MUC16 is maintained within apical membranes of epithelial cells, such as in ocular epithelium^31^, and to be dependent on glycosylation^28^, which may explain why we observe increased apical staining patterns for MUC16 in LGSOC tissues. Beyond its role as a biomarker, MUC16 has been shown to promote cellular proliferation, invasion of ovarian cancer cells^32^, and has previously been shown to discourage immune surveillance, and to be a potential target for immunotherapy^33^. Future efforts will focus on investigating mechanisms of membrane retention of MUC16 in LGSOC and improving our understanding of how this characteristic impacts LGSOC tumor cell biology.

Limitations for this study include small sample sizes for normal fallopian tube, LGSOC, and HGSOC tissues in our discovery cohort. We aimed to address this limitation by prioritizing protein alterations between LGSOC and HGSOC tumors that are also validated within global proteomic data generated for independent cohorts of LGSOC and HGSOC patients. Another limitation may be that samples were obtained by formalin-fixed, paraffin-embedded tissue instead of fresh frozen tissue, however, comparison of fresh tissue samples and fixed frozen paraffin embedded samples has not demonstrated differences in peptide identification and quantification^34^. Strengths of this study include the foundation provided for future proteomic research and the contributions of our findings towards improving our understanding of molecular alterations underlying ovarian cancer pathogenesis.

## Conclusion

Our study contributes to our understanding of global proteome dynamics between normal fallopian tube tissues, LGSOC, and HGSOC tumors, and provides insights into the relative abundance and localization of conserved protein alterations between LGSOC and HGSOC, including CD73 and the ovarian cancer biomarker MUC16/ CA125; these latter findings may also affect clinical interpretation of circulating levels of MUC16/CA125 in these disease subtypes. Future study is needed to determine the significance of protein alterations in the course of this disease, e.g. CD73 and the immune microenvironment of LGSOC tumors, which may serve as additional therapeutic targets for disease treatment.

## Data Availability

Proteomic data generated in this study is accessible at the ProteomeXChange under accession PXD050642.

## Author Contributions

C.M.T., N.W.B., T.P.C. lead the study. C.M.T, P.F., J.O., G.L.M. performed identification of clinical specimens, C.M.T., G.L.M., K.M.D performed clinical data analysis. K.A.C., J.O., and C.M.T. performed sample collections and molecular extraction. N.W.B., J.O., T.A., B.L.H, generated and analyzed proteomics data. P.M.F. P.A.B, S.M.L performed pathology review. C.M.T., N.W.B., wrote the manuscript, P.M.F., J.O., T.A., B.L.H, K.R., K.K.W, D.G., S.L., A.K.S., R.C.B, K.M.D., N.T.P, G.L.M, T.P.C reviewed the manuscript. All authors reviewed and approved the final manuscript.

## Acknowledgements

Funding for this project was provided from the Uniformed Services University of the Health Sciences from the Defense Health Program to the Henry M Jackson Foundation (HJF) for the Advancement of Military Medicine Inc. Gynecologic Cancer Center of Excellence Program, including awards HU0001-20-2-0033, HU0001-21-2-0027, HU0001-22-2-0016, and HU0001-23-2-0038 (PI: Neil T Phippen, Co-PI: G. Larry Maxwell).

## Disclaimer

The views expressed herein are those of the authors and do not reflect the official policy of the Uniformed Services University of the Health Sciences, the Henry M. Jackson Foundation for the Advancement of Military Medicine, Inc., Inova Health System, the Department of Army/Navy/Air Force, Department of Defense, or U.S. Government. Mention of trade names, commercial products, or organizations does not imply endorsement by the U.S. Government.

## Supplementary Tables

1. Supplementary Table 1 (S1): Discovery cohort details
2. Supplementary Table 2 (S2): Global proteome matrix
3. Supplementary Table 3 (S3): Differential analysis of LGSOC vs Fallopian Tube
4. Supplementary Table 4 (S4): Differential analysis of HGSOC vs Fallopian Tube
5. Supplementary Table 5 (S5): Pathway analysis of proteins co-altered between HGSOC and LGSOC vs Fallopian Tube.
6. Supplementary Table 6 (S6): Pathway analysis of proteins uniquely elevated in HGSOC or LGSOC tumors vs Fallopian Tube.
7. Supplementary Table 7 (S7): Differential analysis of LGSOC vs HGSOC
8. Supplementary Table 8 (S8): Proteins significantly co-altered in LGSOC vs HGSOC between Discovery and Validation A & B cohorts.
9. Supplementary Table 9 (S9): Pathway analysis of proteins significantly co-altered between HGSOC and LGSOC tumors in Discovery and Validation A & B cohorts.
10. Supplementary Table 10 (S10): MUC16 immunohistochemistry analysis detailing expert pathology reviews from three independent reviewers, i.e. PATH1, PATH2 and PATH3.

## References

1 Torre, L. A. et al. Ovarian cancer statistics, 2018. CA Cancer J Clin 68, 284–296, doi:10.3322/caac.21456 (2018).

2 Siegel, R. L., Giaquinto, A. N. & Jemal, A. Cancer statistics, 2024. CA Cancer J Clin 74, 12–49, doi:10.3322/caac.21820 (2024).

3 Slomovitz, B. et al. Low-grade serous ovarian cancer: State of the science. Gynecol Oncol 156, 715–725, doi:10.1016/j.ygyno.2019.12.033 (2020).

4 Zarei, S. et al. Clinicopathologic, Immunohistochemical, and Molecular Characteristics of Ovarian Serous Carcinoma With Mixed Morphologic Features of High-grade and Low-grade Serous Carcinoma. Am J Surg Pathol 44, 316–328, doi:10.1097/PAS.0000000000001419 (2020).

5 Gershenson, D. M. et al. Trametinib versus standard of care in patients with recurrent low-grade serous ovarian cancer (GOG 281/LOGS): an international, randomised, open-label, multicentre, phase 2/3 trial. Lancet 399, 541–553, doi:10.1016/S0140-6736(21)02175-9 (2022).

6 Wong, K. K. et al. Integrated multi-omic analysis of low-grade ovarian serous carcinoma collected from short and long-term survivors. J Transl Med 20, 606, doi:10.1186/s12967-022-03820-x (2022).

7 Vallejos, R. et al. Changes in the tumour microenvironment mark the transition from serous borderline tumour to low-grade serous carcinoma. J Pathol, doi:10.1002/path.6338 (2024).

8 Giamougiannis, P., Martin-Hirsch, P. L. & Martin, F. L. The evolving role of MUC16 (CA125) in the transformation of ovarian cells and the progression of neoplasia. Carcinogenesis 42, 327–343, doi:10.1093/carcin/bgab010 (2021).

9 Lee, S. et al. Molecular Analysis of Clinically Defined Subsets of High-Grade Serous Ovarian Cancer. Cell Rep 31, 107502, doi:10.1016/j.celrep.2020.03.066 (2020).

10 Burdett, N. L. et al. Multiomic analysis of homologous recombination-deficient end-stage high-grade serous ovarian cancer. Nat Genet 55, 437–450, doi:10.1038/s41588-023-01320-2 (2023).

11 Ritchie, M. E. et al. limma powers differential expression analyses for RNA-sequencing and microarray studies. Nucleic Acids Res 43, e47, doi:10.1093/nar/gkv007 (2015).

12 Subramanian, A. et al. Gene set enrichment analysis: a knowledge-based approach for interpreting genome-wide expression profiles. Proc Natl Acad Sci U S A 102, 15545–15550, doi:10.1073/pnas.0506580102 (2005).

13 Tang, D. et al. SRplot: A free online platform for data visualization and graphing. PLoS One 18, e0294236, doi:10.1371/journal.pone.0294236 (2023).

14 Dinh, H. Q. et al. Single-cell transcriptomics identifies gene expression networks driving differentiation and tumorigenesis in the human fallopian tube. Cell Rep 35, 108978, doi:10.1016/j.celrep.2021.108978 (2021).

15 Ghosh, A., Syed, S. M. & Tanwar, P. S. In vivo genetic cell lineage tracing reveals that oviductal secretory cells self-renew and give rise to ciliated cells. Development 144, 3031–3041, doi:10.1242/dev.149989 (2017).

16 Yan, S. et al. The Potential of Targeting Ribosome Biogenesis in High-Grade Serous Ovarian Cancer. Int J Mol Sci 18, doi:10.3390/ijms18010210 (2017).

17 Kang, J. et al. Ribosomal proteins and human diseases: molecular mechanisms and targeted therapy. Signal Transduct Target Ther 6, 323, doi:10.1038/s41392-021-00728-8 (2021).

18 Lopez-Reig, R. & Lopez-Guerrero, J. A. The hallmarks of ovarian cancer: proliferation and cell growth. EJC Suppl 15, 27–37, doi:10.1016/j.ejcsup.2019.12.001 (2020).

19 Casey, R. C. & Skubitz, A. P. CD44 and beta1 integrins mediate ovarian carcinoma cell migration toward extracellular matrix proteins. Clin Exp Metastasis 18, 67–75, doi:10.1023/a:1026519016213 (2000).

20 Yi, B. R., Kim, T. H., Kim, Y. S. & Choi, K. C. Alteration of epithelial-mesenchymal transition markers in human normal ovaries and neoplastic ovarian cancers. Int J Oncol 46, 272–280, doi:10.3892/ijo.2014.2695 (2015).

21 Ploeg, E. M. et al. A Novel Bispecific Antibody for EpCAM-Directed Inhibition of the CD73/Adenosine Immune Checkpoint in Ovarian Cancer. Cancers (Basel) 15, doi:10.3390/cancers15143651 (2023).

22 Mirza, M. R. et al. NSGO-OV-UMB1/ENGOT-OV30: A phase II study of durvalumab in combination with the anti-CD73 monoclonal antibody Oleclumab in patients with relapsed ovarian cancer. Gynecol Oncol 188, 103–110, doi:10.1016/j.ygyno.2024.06.017 (2024).

23 Turcotte, M. et al. CD73 is associated with poor prognosis in high-grade serous ovarian cancer. Cancer Res 75, 4494–4503, doi:10.1158/0008-5472.CAN-14-3569 (2015).

24 Wong, K.-K. & Gershenson, D. M. Abstract 135: Immunosuppressive microenvironment in low-grade serous ovarian carcinoma. Cancer Research 78, 135–135, doi:10.1158/1538-7445.am2018-135 (2018).

25 Yue, T. et al. Two Similar Signatures for Predicting the Prognosis and Immunotherapy Efficacy of Stomach Adenocarcinoma Patients. Front Cell Dev Biol 9, 704242, doi:10.3389/fcell.2021.704242 (2021).

26 Das, S. et al. Membrane proximal ectodomain cleavage of MUC16 occurs in the acidifying Golgi/post-Golgi compartments. Sci Rep 5, 9759, doi:10.1038/srep09759 (2015).

27 Bruney, L., Conley, K. C., Moss, N. M., Liu, Y. & Stack, M. S. Membrane-type I matrix metalloproteinase-dependent ectodomain shedding of mucin16/ CA-125 on ovarian cancer cells modulates adhesion and invasion of peritoneal mesothelium. Biol Chem 395, 1221–1231, doi:10.1515/hsz-2014-0155 (2014).

28 Taniguchi, T. et al. N-Glycosylation affects the stability and barrier function of the MUC16 mucin. J Biol Chem 292, 11079–11090, doi:10.1074/jbc.M116.770123 (2017).

29 But, I. & Gorisek, B. Preoperative value of CA 125 as a reflection of tumor grade in epithelial ovarian cancer. Gynecol Oncol 63, 166–172, doi:10.1006/gyno.1996.0301 (1996).

30 Wang, Y. et al. MUC16 expression during embryogenesis, in adult tissues, and ovarian cancer in the mouse. Differentiation 76, 1081–1092, doi:10.1111/j.1432-0436.2008.00295.x (2008).

31 Mantelli, F. & Argueso, P. Functions of ocular surface mucins in health and disease. Curr Opin Allergy Clin Immunol 8, 477–483, doi:10.1097/ACI.0b013e32830e6b04 (2008).

32 Felder, M. et al. MUC16 (CA125): tumor biomarker to cancer therapy, a work in progress. Mol Cancer 13, 129, doi:10.1186/1476-4598-13-129 (2014).

33 Zhai, Y. et al. MUC16 affects the biological functions of ovarian cancer cells and induces an antitumor immune response by activating dendritic cells. Ann Transl Med 8, 1494, doi:10.21037/atm-20-6388 (2020).

34 Bateman, N. W. & Conrads, T. P. Recent advances and opportunities in proteomic analyses of tumour heterogeneity. J Pathol 244, 628–637, doi:10.1002/path.5036 (2018).

